# Declaring a tuberculosis outbreak over with genomic epidemiology

**DOI:** 10.1101/040527

**Authors:** Hollie-Ann Hatherell, Xavier Didelot, Sue L. Pollock, Patrick Tang, Anamaria Crisan, James C. Johnston, Caroline Colijn, Jennifer Gardy

## Abstract

We report an updated method for inferring the time at which an infectious disease was transmitted between persons from a time-labelled pathogen genome phylogeny. We applied the method to 48 Mycobacterium tuberculosis genomes as part of a real-time public health outbreak investigation, demonstrating that although active tuberculosis (TB) cases were diagnosed through 2013, no transmission events took place beyond mid-2012. Subsequent cases were the result of progression from latent TB infection to active disease and not recent transmission. This evolutionary genomic approach was used to declare the outbreak over in January 2015.

## Introduction

Genomics is revolutionizing public health practice (Kwong et al., 2015). Mutational and evolutionary events within a pathogen population not only have consequences for the disease, but also present opportunities for understanding transmission and developing targeted public health interventions. Inferring person-to-person transmission from genomic data is one such example – genome sequencing has now helped identify individual infection events in multiple outbreaks at levels from hospital wards to communities to countries (Croucher & Didelot, 2015).

Transmission inference from genomic data uses mutations – fixed or minor variants (Worby et al., 2014; Poon et al., 2016) – shared across outbreak isolates to identify putative infection events. We previously developed TransPhylo (Didelot et al., 2014) – a Bayesian method for inferring transmissions and their timing given mutational events captured in a time-labelled phylogeny – and used it to reconstruct transmissions between the first 33 cases (2008-2011) of a large TB outbreak. The outbreak in question began with the May 2008 diagnosis of a highly infectious client in a homeless shelter in British Columbia, Canada and peaked in 2010. Intensive case-finding in the community ultimately screened 2,310 individuals and a total of 52 TB cases were diagnosed through December 2013 (Figure 1A) (Cheng et al., 2015).

**Figure 1.**
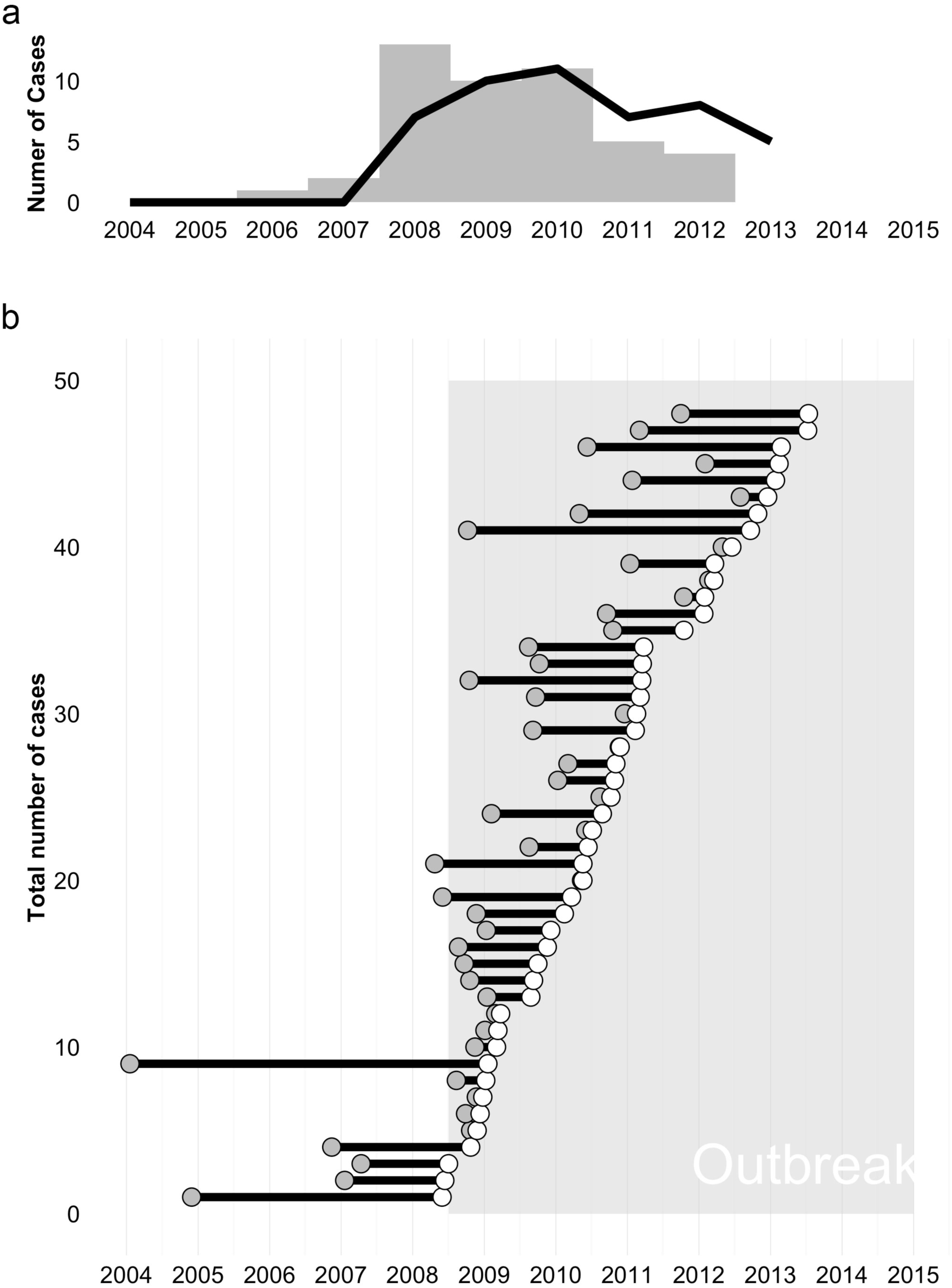
Timing of infections inferred from genomic data. Panel A: The outbreak’s epidemic curve based on time of diagnosis (black line) or Tinf – the time of infection estimated by TransPhylo (grey bars). Panel B: Each case’s infected period is shown as a line originating at Tinf (grey dot) and continuing until the case was diagnosed (white dot). The outbreak period (May 2008-January 2015) is indicated with shading.

In the absence of a formal definition, a TB outbreak is generally deemed over when transmission of the outbreak strain has stopped for >2 years; however, latent TB infection (LTBI) complicates declaring an outbreak’s end. Amongst individuals identified with LTBI, only 5-10% will progress to active disease, with most developing disease within two years of infection (WHO, 2015); however, delayed progression occurring more than two years after infection is not uncommon. As incident case numbers begin to decline, TB controllers must differentiate LTBI cases acquired >2 years ago and only now progressing to active disease from new cases that were recently acquired, suggesting ongoing transmission.

One of TransPhylo’s outputs is *T*_*inf*_ – the estimated time at which an individual was infected. *T*_*inf*_ can differentiate delayed progression from new infection; however, TransPhylo’s underlying SIR model – and indeed all compartmental epidemic models – does not capture the true size of the susceptible population or the true variation in the infectious periods, which may affect inferred *T*_*inf*_ values.

In September 2014, at the request of the Medical Health Officer leading the outbreak response, we analysed Mycobacterium tuberculosis genomes from 48 of the 52 cases to determine whether a decline in newly diagnosed cases truly signaled the outbreak’s end. We replaced TransPhylo’s SIR model with a branching model to better infer the timing of transmission amongst the 48 cases and asked whether cases diagnosed in 2013 were the result of recent transmission or delayed progression of an infection acquired earlier in the outbreak.

## Methods

As part of an earlier investigation during the outbreak, we sequenced 33 M. tuberculosis genomes from outbreak cases diagnosed between May 2008 and April 2011 (Didelot et al., 2014). In September 2014, we sequenced genomes from a further 15 cases on the MiSeq platform, for a total of 48 genomes (Accession: PRJEB12764, Data Citation 1). Four of the 52 outbreak cases did not have an M. tuberculosis isolate available for sequencing as they were diagnosed out-of-province or on clinical grounds during a post-mortem.

Reads were mapped against the M. tuberculosis CDC1551 reference genome (NC_002755.2) using BWAmem (Li & Durbin, 2009) and variants called using samtools mpileup (Li et al., 2009); the commands we used are available in a text file from FigShare (DOI: 10.6084/m9.figshare.2077390, Data Citation 2).

From the resulting VCF files, we removed all variant positions that were identical across all 48 genomes, leaving only variants that differentiate outbreak isolates. We filtered a matrix of these positions to remove variant positions within 150bp of another variant position, suggesting misalignment to a low-complexity region, as well as variant positions without an mpileup QUAL score equal to 222 in at least one isolate. This left 28 positions across the 48 isolates, which were manually reviewed before further analysis; a FASTA file of these variants labeled with isolate sampling date (in the format “days since X”) is available at FigShare (DOI: 10.6084/m9.figshare.2077405, Data Citation 3). Variants were concatenated and analysed with BEAST (Drummond & Rambaut, 2007) and the resulting timed phylogeny was passed to TransPhylo for transmission timing inference.

We used a modified version of TransPhylo (github.com/xavierdidelot/TransPhylo, Data Citation 4) in which we replaced the existing SIR epidemic model with a branching model. The mathematics describing the branching model are detailed in a Methods Supplement included here.

## Results

Examining *T*_*inf*_ for each of the 48 sequenced cases revealed that the last person-to-person transmission occurred in late July or early August 2012 (Figure 1A, B). The eight cases diagnosed afterwards were all instances of delayed progression, with most of the transmission events leading to these cases occurring in 2010-2011. Indeed, the epidemic curves based on *T*_*inf*_ and on diagnosis date (Figure 1A) show echo each other, with transmission concentrated largely before 2011 and diagnoses extending two years beyond that, underscoring the importance of early active case-finding and preventive therapy in a TB outbreak.

The average time from infection to diagnosis was 1.2 years (95%CI ± 0.31; Figure 1B), increasing from 0.98 (95%CI ± 0.82) in 2009 to 1.97 (95%CI ± 0.69) in 2013 despite continued intensive surveillance. This supports the hypothesis that later cases were largely due to delayed progression of infection acquired earlier in the outbreak and highlights the need for extensive follow-up of infected contacts and provision of LTBI preventive therapy.

## Conclusion

We presented our findings to the Medical Health Officer and the Outbreak Management Team on January 9, 2015. After considering our genomic evidence indicating that no transmission of the outbreak strain had been detected since 2012 – thereby fulfilling the criteria for at least two years without a transmission event – and corroborating evidence from the ongoing epidemiological investigation, the outbreak was declared over on January 29, 2015 (Interior Health Authority, 2015).

This is the first demonstration that evolutionary genomic analysis can be used to declare a complex community outbreak over, suggesting a new role for public health genomics in not just identifying transmission events, but also in timing these events to better understand an outbreak’s dynamics and guide the realtime public health response.

## Acknowledgments

This work was supported by the Engineering and Physical Sciences Research Council (H.H, C.C – grant EP/K026003/1), the Canada Research Chairs program (J.L.G), and the Michael Smith Foundation for Health Research (J.C.J). We thank the members of the Interior Health Authority TB Outbreak Management Team, the BCCDC Public Health Laboratory’s Mycobacteriology Laboratory, and BCCDC TB Services for their assistance, particularly Lori Hiscoe, Rob Parker, Clare Kong, Mabel Rodrigues, and Victoria Cook.

## Data Bibliography

1. Gardy, J. European Nucleotide Archive http://www.ebi.ac.uk/ena/data/view/PRJEB12764 (2016).

2. Gardy, J. FigShare http://dx.doi.org/10.6084/m9.figshare.2077390 (2016).

3. Gardy, J. FigShare http://dx.doi.org/10.6084/m9.figshare.2077405 (2016)

4. Didelot, X. GitHub https://github.com/xavierdidelot/TransPhylo) (2015).

## References

Cheng, J.M., Hiscoe, L., Pollock, S.L., Hasselback, P., Gardy, J.L., Parker, R. (2015). A clonal outbreak of tuberculosis in a homeless population in the interior of British Columbia, Canada, 2008-2015. Epidemiol Infect. 143, 3220–3226.

Croucher, N.J., Didelot, X. (2015). The application of genomics to tracing bacterial pathogen transmission. Curr Opin Microbiol. 23, 62–67.

Didelot, X., Gardy, J., Colijn, C. (2014). Bayesian inference of infectious disease transmission from whole-genome sequence data. Mol Biol Evol. 31, 869–1879.

Interior Health Authority. (2015). Six year TB outbreak comes to an end. https://www.interiorhealth.ca/AboutUs/MediaCentre/NewsReleases/Documents/Six%20year%20TB%20outbreak%20comes%20to%20an%20end.pdf, last accessed January 14, 2016.

Kwong, J.C, McCallum, N., Sintchenko, V., Howden, B.P. (2015). Whole genome sequencing in clinical and public health microbiology. Pathology. 47, 199–210.

Poon, L.L., Song, T., Rosenfeld, R., Lin, X., Rogers, M.B., Zhou, B., Sebra, R., Halpin, R.A., Guan, Y. & other authors. (2016). Quantifying influenza virus diversity and transmission in humans. Nat Genet. doi: 10.1038/ng.3479.

Worby, C.J., Chang, H.H., Hanage, W.P., Lipsitch, M. (2014). The distribution of pairwise genetic distances: a tool for investigating disease transmission. Genetics. 198, 1395–1404.

World Health Organization. (2015). Guidelines on the Management of Latent Tuberculosis Infection. http://apps.who.int/iris/bitstream/10665/136471/1/9789241548908_eng.pdf?ua=1&ua=1, last accessed January 14, 2016.

